# The effect of acute sleep deprivation on skeletal muscle protein synthesis and the hormonal environment

**DOI:** 10.1101/2020.03.09.984666

**Authors:** Séverine Lamon, Aimee Morabito, Emily Arentson-Lantz, Olivia Knowles, Grace Elizabeth Vincent, Dominique Condo, Sarah Alexander, Andrew Garnham, Douglas Paddon-Jones, Brad Aisbett

**Author notes:** Corresponding author: Séverine Lamon, PhD Institute for Physical Activity and Nutrition Research School of Exercise and Nutrition Sciences Deakin University, 221 Burwood Hwy, Burwood 3125. Australia.

## Abstract

Chronic sleep loss is a potent catabolic stressor, increasing the risk of metabolic dysfunction and loss of muscle mass and function. To provide mechanistic insight into these clinical outcomes, we sought to determine if acute sleep deprivation blunts skeletal muscle protein synthesis and promotes a catabolic environment. Healthy young adults (N=13; 7 male, 6 female) were subjected to one night of total sleep deprivation (DEP) and normal sleep (CON) in a randomized cross-over design. Anabolic and catabolic hormonal profiles, skeletal muscle fractional synthesis rate and markers of muscle protein degradation were assessed across the following day. Acute sleep deprivation reduced muscle protein synthesis by 18% (CON: 0.072 ± 0.015 vs. DEP: 0.059 ± 0.014 %•h^-1^, *p=0.040*). In addition, it increased plasma cortisol by 21% (*p=0.030*) and decreased plasma testosterone, but not IGF-1, by 22% (*p=0.029*). A single night of total sleep deprivation is sufficient to induce anabolic resistance and a pro-catabolic environment. These acute changes may represent mechanistic precursors driving the metabolic dysfunction and body composition changes associated with chronic sleep deprivation.

## Introduction

Acute and chronic sleep loss are linked with a range of negative physiological and psychological outcomes (22). While complete sleep deprivation rapidly impedes simple and complex cognitive functions, sleep restriction impairs whole-body homeostasis, leading to undesirable metabolic consequences in the short- and longer-term (39). Most metabolic tissues including liver, adipose tissue and skeletal muscle are at risk of developing sleep loss-associated adverse outcomes.

Skeletal muscle is a primary regulator of human metabolism. Sleep deprivation (9, 10) and restriction (19) have the potential to profoundly affect muscle health by altering gene regulation and substrate metabolism. Even relatively short periods of sleep restriction (less than a week) can compromise glucose metabolism, reduce insulin sensitivity and impair muscle function (5, 7). Skeletal muscle is made up of 80% of proteins and maintaining optimal muscle protein metabolism is equally critical for muscle health. In situations where skeletal muscle protein synthesis chronically lags protein degradation, a loss of muscle mass is inevitable. Low muscle mass is a hallmark of and precursor to a range of chronic health conditions, including neuromuscular disease, sarcopenia and frailty, obesity and type II diabetes (41). Population-based studies report that the risk of developing these conditions is 15-30% higher in individuals who regularly experience sleep deprivation, sleep restriction, and inverted sleep-wake cycles (26, 31, 50). To this end, a growing body of evidence suggests that a lack of sleep may directly affect muscle protein metabolism (1, 35, 42).

Rodent studies first demonstrated a possible causal link between complete sleep deprivation and disrupted muscle protein metabolism. Rats subjected to 96 h of paradoxical sleep deprivation, where rapid eye movement sleep is restricted, experienced a decrease in muscle mass (13) and muscle fibre cross-sectional area (15). In this model, sleep deprivation negatively impacted the pathways regulating protein synthesis and increased muscle proteolytic activity (15). These findings were paralleled by a human study reporting a catabolic gene signature in skeletal muscle following one night of total sleep deprivation in healthy young males (10). To expand on this acute model, investigators recently demonstrated that five consecutive nights of sleep restriction (four hours per night) reduced myofibrillar protein synthesis in healthy young males when compared to normal sleep patterns (42). The possible mechanisms underlying these effects have not been investigated, but might involve the hormonal environment.

Factors that regulate skeletal muscle protein metabolism at the molecular level are influenced by mechanical (muscle contraction), nutritional (dietary protein intake) and hormonal inputs (41). Testosterone and IGF-1 positively regulate muscle protein anabolism by promoting muscle protein synthesis (43, 46), while repressing the genes that activate muscle protein degradation (51). In contrast, cortisol drives catabolism by activating key muscle protein degradation pathways (21). Experimental evidence suggests that acute and chronic sleep loss alters anabolic (29, 40) and catabolic (10, 14) hormone secretion patterns in humans, providing a possible mechanism for impaired muscle protein metabolism.

While our understanding of the health consequences of sleep deprivation continues to improve, important gaps and opportunities remain. This includes linking acute mechanistic changes with clinically observable outcomes and moving towards a more prescriptive, individualized understanding of sleep deprivation by examining sex-based differences. In this study, we sought to determine if one night of complete sleep deprivation promotes a catabolic hormonal environment and compromises post-prandial muscle protein synthesis and markers of muscle degradation in young, healthy male and female participants.

## Methods

### Participants

Thirteen young (18-35 years old), healthy male and female students gave their informed consent to participate in this randomized, crossover-designed study. Participants were excluded if they had a history of recent transmeridian travel (i.e., no travel across multiple time zones in the previous four weeks), shiftwork (i.e., no involvement in shiftwork over the previous three months), frequent napping (i.e., ≥ 2 naps per week), or had a diagnosed sleep disorder. Participants were required to have habitual bed (2200–0000) and wake (0600–0800) times that were broadly consistent with the experimental protocol and to self-report obtaining a minimum of 7 hours of sleep (not time in bed) per night. Chronotype was assessed using the morningness-eveningness (ME) questionnaire (20). Participants exhibiting extreme morningness (score > 70) or eveningness (score < 30) were excluded. All participants but three displayed an ‘intermediate’ ME type. A detailed account of the strategy for female volunteer recruitment and testing have been comprehensively described by our group (24). Briefly, effects of female reproductive hormone fluctuations were minimised by testing all female participants during the same phase of their menstrual cycle in both conditions. Although it has previously been shown that the menstrual cycle had no effect on female muscle protein synthesis (34), our primary outcome, the follicular phase was avoided to ensure the ratio of estrogen to progesterone was at its lowest. The study was approved by the Deakin University Human Research Ethics Committee (2016-028) and conducted in accordance to *The Declaration of Helsinki* (1964) and its later amendments. Participants’ physiological characteristics, ME score, and self-reported habitual time asleep are summarized in Table 1. There were no sex-specific difference in ME score (*p=0.148*) or self-reported habitual time asleep (*p=0.401*).

**Table 1.** Participants characteristics. Mean ± SD; BMI = Body Mass Index; ME = Morningness Eveningness score

| Sex | Age | Mass (kg) | Height<br>(cm) | BMI<br>(kg·m<sup>-2</sup>) | ME score | Habitual time<br>asleep (self-reported) (h) |
| --- | --- | --- | --- | --- | --- | --- |
| Male (N=7) | 22 ± 1.8 | 71.6 ± 11.3 | 173.5 ± 9.0 | 22.6 ± 4.1 | 48.0 ± 6.4 | 7.5 ± 0.6 |
| Female (N=6) | 20 ± 1.3 | 60.1 ± 10.3 | 170.5 ± 5.1 | 20.7 ± 3.2 | 53.8 ± 6.6 | 7.8 ± 0.7 |

### Sample size calculation

At the time this study was designed, there was no published study investigating the effect of sleep deprivation on muscle protein synthesis. Using data from studies investigating the effects of an anabolic and catabolic stimulus (e.g., immobilization or exercise) on changes in muscle protein synthesis (28, 36), power analyses conducted on our primary outcome (fractional synthesis rate) indicated that a sample size of 13 would minimize the risk of type II error (β =0.2, α =0.05). Males and females were included as previous work demonstrated that muscle fractional synthesis rate, our primary outcome, is similar in both sexes (24, 49).

### Pre-study procedure

During the week priot to the study, participants were instructed to maintain their habitual sleep behaviour. Participants wore an actigraph (Actical MiniMitter/Respironics, Bend, OR) on their non-dominant wrist, and completed a sleep diary. The diary was used to corroborate actigraphy data and minimise possibility of incorrectly scoring periods of sedentary wakefulness as sleep.

Participants completed a control (CON) and experimental (DEP; sleep deprivation) trial in a randomized, crossover design. Trials were separated by at least four weeks to allow for full recovery. Forty-eight hours prior to each trial, participants were required to refrain from strenuous exercise, alcohol and caffeine. On the night of the trial (CON or DEP), a standardized meal containing approximately 20% fat, 14% protein and 66% carbohydrate (energy intake ranging between 8.4-8.9 kcal/kg) was provided to participants with water ad libitum.

### Study procedure

On the night of the sleep deprivation trial (DEP), participants consumed a standardized meal at 1900 and reported to the laboratory at 2100 where they were limited to sedentary activities (i.e., reading a book, watching a movie). Participants were constantly observed by research personnel and monitored by actigraphy to ensure they did not fall asleep. They remained in a sound attenuated, light (211±14 lux) and temperature (21±2°C) controlled facility for the entire 30h protocol. Participants were permitted to consume low-protein snacks (i.e., fruits and vegetables) and water *ad libitum* during the sleep deprivation period. Regardless of potential differences in insulinemia, adding a non-pharmacological dose of carbohydrates to a protein synthesis activating dose of proteins (15-30 g) has no additive effect on fractional synthesis rate (17, 18, 25, 44), our primary outcome. For the control trial (CON), participants consumed a standardized meal at 1900 and were permitted to sleep from 2200 to 0700 at home, rather than risking a night of disrupted sleep in an unfamiliar laboratory/clinical environment. At 0700 the following morning, a researcher and nurse with pre-arranged access to the participants’ home woke the participant and immediately collected a venous blood sample prior to any physical activity or light exposure. Participants were then transported to the laboratory to complete the experimental protocol.

For both DEP and CON trials, at 0730 participants consumed a standardized breakfast containing approximately 9% fat, 11% proteins and 80% carbohydrates, and 20.3 ± 1.8 g of proteins. Rather than fasting our participants, standardized meals were provided as part of the experimental protocol for DEP and CON. The goal was to: i) reduce participant discomfort and improve compliance, and ii) model a more realistic and balanced, post-prandial metabolic environment, rather than the overtly catabolic environment associated with 24h of fasting. At 0800, an 18-gauge cannulae was inserted into the antecubital vein of each arm for blood sampling and the primed (0.34 mg•kg^-1^), constant infusion (0.0085 mg•kg^-1^•min^-1^) of [ring-^13^C_6_]-L-phenylalanine (Cambridge Isotope Laboratories, Tewksbury, MA) from 1000 to the end of the protocol. At 1200, participants consumed a standardized lunch containing 12% fat, 21% protein and 67% carbohydrate, and 20.6 ± 0.3 g protein. The slowly digested, whole-food meals reduced the fluctuations in plasma Phe enrichment, thus avoiding the need to add tracer to the meals (33). Skeletal muscle samples were obtained at 1300 and 1500 under local anaesthesia (1% Lidocaine) at separate locations from the belly of the *vastus lateralis* muscle using a percutaneous needle biopsy technique as previously described by our group (28). Muscle samples were immediately frozen in liquid nitrogen and used for the measurement of isotopic enrichment and gene expression analysis. An outline of the experimental protocol is presented in Figure 1.

**Figure 1.**
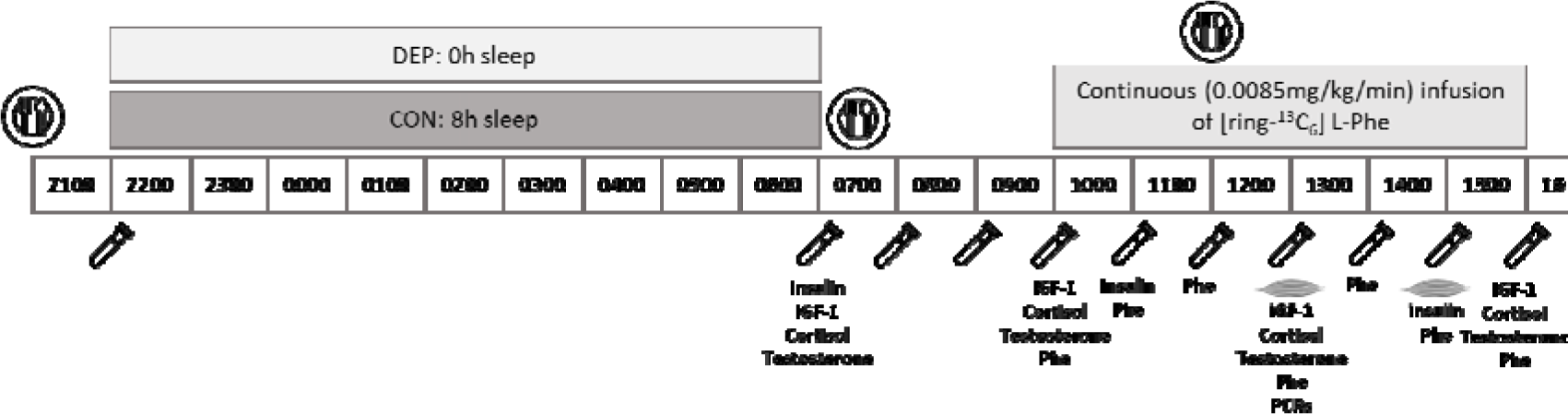
Experimental 178 Protocol. 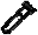 blood collection; 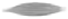: muscle collection; 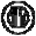: standardized meal. IGF-1, cortisol and testosterone concentrations were measured at the 0700, 1000, 1300 and 1600 timepoints. Insulin concentrations were measured at the 0700, 1100 and 1500 timepoints. Phe enrichment (Phe) was measured in both muscle tissue and blood samples between 1000 and 1600. PCRs were run on muscle tissue collected at 1300.

### Sleep measures

Sleep was recorded objectively using actigraphy (Actical MiniMitter/Respironics, Bend, OR). The Actical (28 × 27 × 10 mm, 17 g) device uses a piezoelectric omnidirectional accelerometer, which is sensitive to movements in all planes in the range of 0.5–3.0 Hz. Data output from activity monitors (actigraphy) provides an objective, non-invasive, indirect assessment of sleep and has been validated against polysomnography (2). Primary outcomes were total sleep time and sleep efficiency (total sleep time/time in bed).

### Hormone measures

Venous blood samples were collected every hour from 0700 to 1700 in EDTA-tubes, manually inverted and immediately centrifuged for 15 min at 13,000 rev•min^-1^ at 4°C. The supernatant (plasma) was then isolated and frozen at -80°C for further analysis. Plasma cortisol, testosterone and insulin growth factor-1 (IGF-1) concentrations were determined using a highLJsensitivity enzyme immunoassay ELISA kit (IBL International, Hamburg, Germany) according to the manufacturer s instructions. Insulin concentration was determined using the MILLIPLEX^®^ MAP Human Metabolic Hormone Magnetic Bead Panel (Merck KGaA, Darmstadt, Germany) according to the manufacturer s instructions.

### Isotopic enrichment in plasma

After thawing, plasma was precipitated using an equal volume of 15% sulfosalicylic acid (SSA) solution and centrifuged for 20 min at 13,000 rev•min^-1^ at 4°C. Blood amino acids were extracted from 500 uL of supernatant by cation exchange chromatography (Dowex AG 50W-8X, 100–200 mesh H+ form; Bio-Rad Laboratories). Phenylalanine enrichments were determined by gas chromatography–mass spectrometry (GC-MS) using the tertbutyldimethylsilyl derivative with electron impact ionization as described previously (16). Ions 336 and 342 were monitored.

### Isotopic enrichment in muscle proteins

A 30 mg piece of muscle was used for isolation of mixed muscle bound and intracellular protein fractions. Briefly, bound muscle proteins were extracted in perchloric acid and hydrolysed using 6N hydrochloric acid (110°C for 24 h). Isotopic enrichments of [ring-_13_C^6^]-L-phenylalanine in tissue fluid (intracellular fraction) were used as a precursor pool for the calculation of the fractional synthesis rate. Total muscle phenylalanine was isolated using cation exchange chromatography (50W-8X, 200–400 mesh H+ form; Bio-Rad Laboratories). Amino acids were eluted in 8 mL of 2N ammonium hydroxide and dried under vacuum. Muscle intracellular and bound protein [ring-_13_C^6^]-L-phenylalanine enrichments were determined by GC-MS with electron impact ionization using the tert-butyldimethylsilyl derivative. Ions 238 and 240 were monitored for bound protein enrichments; ions 336 and 342 were monitored for intracellular enrichments as described previously (16). Mixed muscle protein FSR (% / hour) was calculated by measuring the direct incorporation of [ring-_13_C^6^]-L-phenylalanine by using the precursor-product model (36):

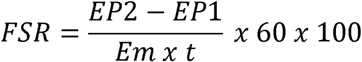

where EP1 and EP2 are the bound enrichments of [ring-_13_C^6^]-L-phenylalanine for the 2 muscle biopsies, Em is the mean enrichment of [ring-_13_C^6^]-L-phenylalanine in the muscle intracellular pool, and t is the time interval (min) between biopsies.

### RNA extraction and gene expression analysis

Muscle biopsies collected at 1300 were used for gene expression analysis. RNA was extracted from ~15 mg of skeletal muscle samples using Tri-Reagent© Solution (Ambion Inc., Austin, TX, USA) according to the manufacturer s protocol. RNA was treated with DNase I Amplification Grade (Thermo Fisher Scientific, MA) and RNA concentration was assessed using the Nanodrop 1000 Spectrophotometer (Thermo Fisher Scientific). First-strand cDNA was generated from 1000 ng RNA using the High Capacity RT-kit (Applied Biosystems, Carlsbad, CA, USA). cDNA was then treated with RNase H (Thermo Fisher Scientific) according to the manufacturer protocol. Real-time PCR was carried out using an AriaMx real-time PCR system (Agilent Technologies, Santa Clara, CA) to measure mRNA levels. mRNA levels for ARNTL (BMAL1), CRY1, PER1, IGF-1Ea, IGF-1Eb, FBX032 (atrogin-1), TRIM63 (MuRF-1), FOXO1 and FOXO3 were measured using 1 × SYBR© Green PCR MasterMix (Applied Biosystems) and 5 ng of cDNA. All primers were used at a final concentration of 300 nM. Primer details are provided in Table 2. Single-strand DNA was quantified using the Quant it OliGreen ssDNAAssay Kit (Thermo Fisher Scientific) according to the manufacturer s instruction. ssDNA was used for PCR normalization as previously validated in (32). No differences in ssDNA concentrations were found between groups (data can be found at https://doi.org/10.6084/m9.figshare.12629972.v1). This normalization strategy was cross-checked against the common housekeeper gene GAPDH (data not shown).

**Table 2.**
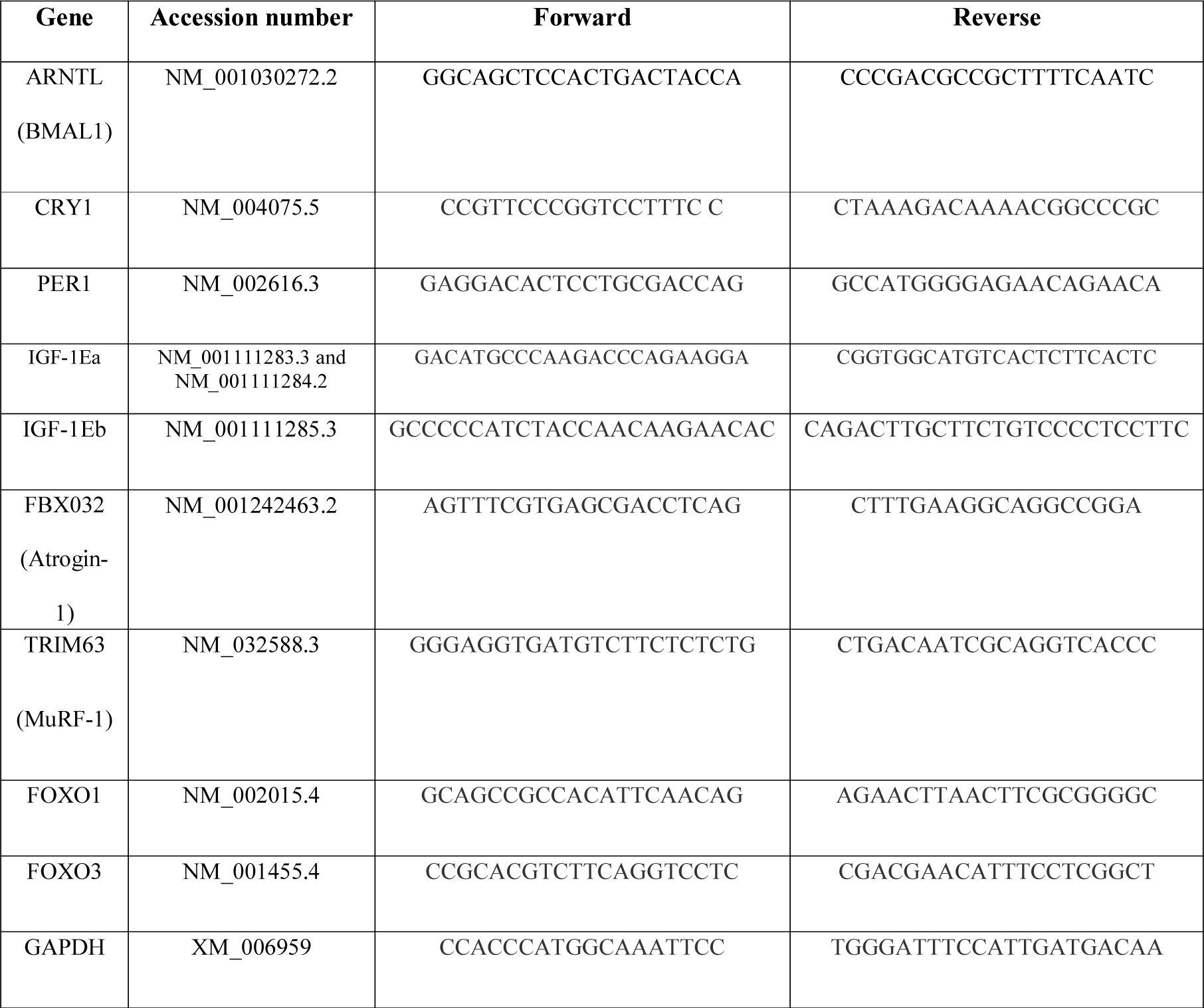
Primer sequences

### Statistical analysis

Statistical analyses were conducted using SPSS 26.0 (IBM Corp, Armonk, NY). Diagnostic plots of residuals and fitted values were checked to ensure homogeneity of the variance. Hormonal levels were analysed using a two-way analysis of variance (ANOVA) with within-participant factors for time and condition (CON vs DEP) unless specified otherwise. The Sidak test was used to compare pairs of means when a main effect was identified. For FSR and hormone concentrations, single-tailed paired t-tests were used to compare group means. For gene expression data, two-tailed paired t-tests were used to compare group means. Area under the curve (AUC) was computed on hormone values using the trapezoidal method. The significance levels for the F-tests in the t-tests and ANOVA and the Sidak tests were set at p<0.05. All data are reported as mean ± SD.

## Results

### Sleep

During the week prior to the study, there were no differences in total sleep time (CON: 5.9 ± 0.5 h, DEP: 6.1 ± 1.4 h, *p = 0.718*) or sleep efficiency (CON: 78.5 ± 6.5 %, DEP: 79.4 ± 4.7 %, *p = 0.801*). Similarly, during the night directly preceding the sleep intervention, there were no differences in total sleep time (CON: 6.8 ± 0.8 h, DEP: 7.4 ± 0.7 h, *p = 0.195*) or sleep efficiency (CON: 77.3 ± 6.3 %, DEP: 81.0 ± 8.6 %, *p = 0.424*).

### Muscle protein synthesis

Subjects remained in isotopic steady state for the duration of the isotope infusion, with no differences in plasma enrichment between CON and DEP conditions (Figure 2A). Sleep deprivation reduced post-prandial muscle protein fractional synthesis rate (FSR) by 18% (CON: 0.072 ± 0.015 vs. DEP: 0.059 ± 0.014 %•h^-1^, *p=0.040*) (Figure 2B).

**Figure 2.**
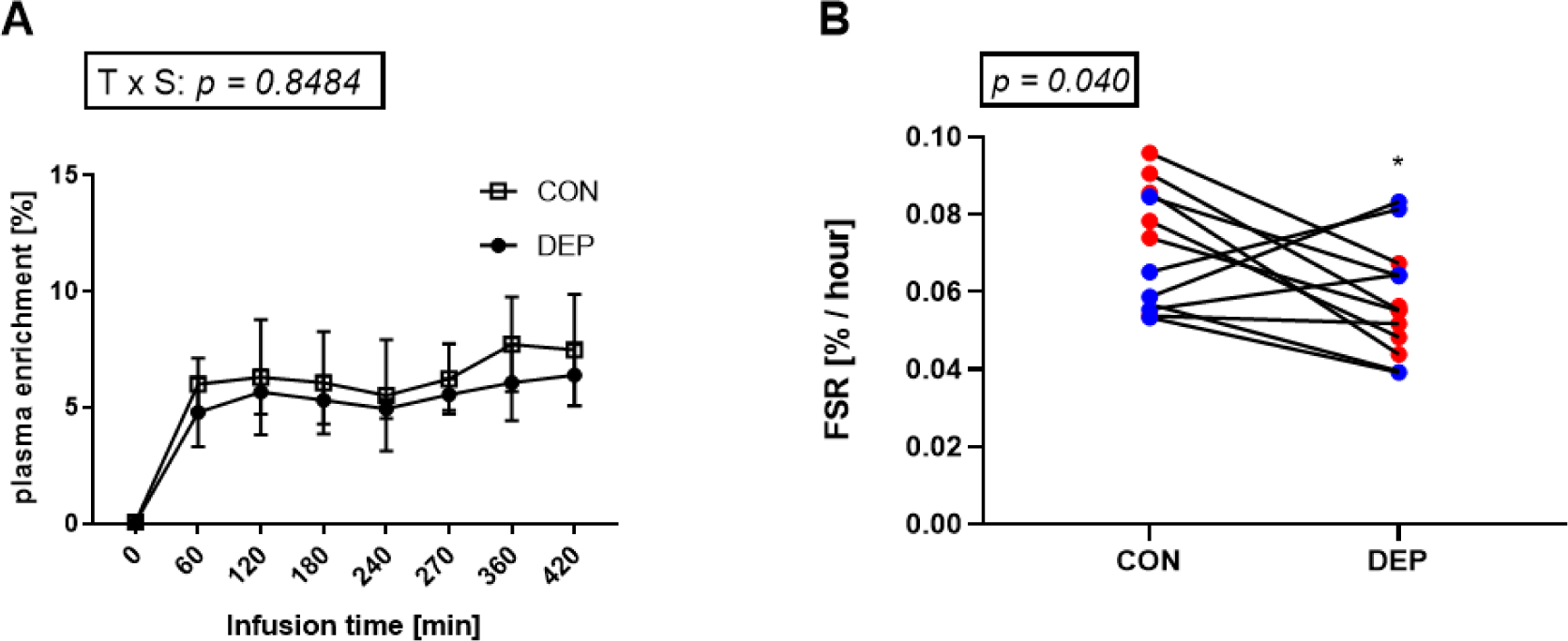
Plasma enrichment of [ring-^13^C_6_]-L-phenylalanine during the experimental protocol (N=4). Data are presented as mean ± SD (A). Post-prandial mixed muscle fractional synthesis rate measured in the control (CON) and sleep-deprived (DEP) conditions. Red dots depict male subjects. Blue dots depict female subjects (B).

### Plasma testosterone levels

There was a main effect of time (*p=0.002*) but the interaction effect of sleep × time for plasma testosterone levels did not reach statistical significance (*p=0.063*; Figure 3A). The area under the curve decreased by 22% in the DEP condition (*p=0.029*; Figure 3B). A male and a female sub-population group were visually highlighted, where all male subjects had their testosterone AUC decreasing in the DEP condition (Figure 3B).

**Figure 3.**
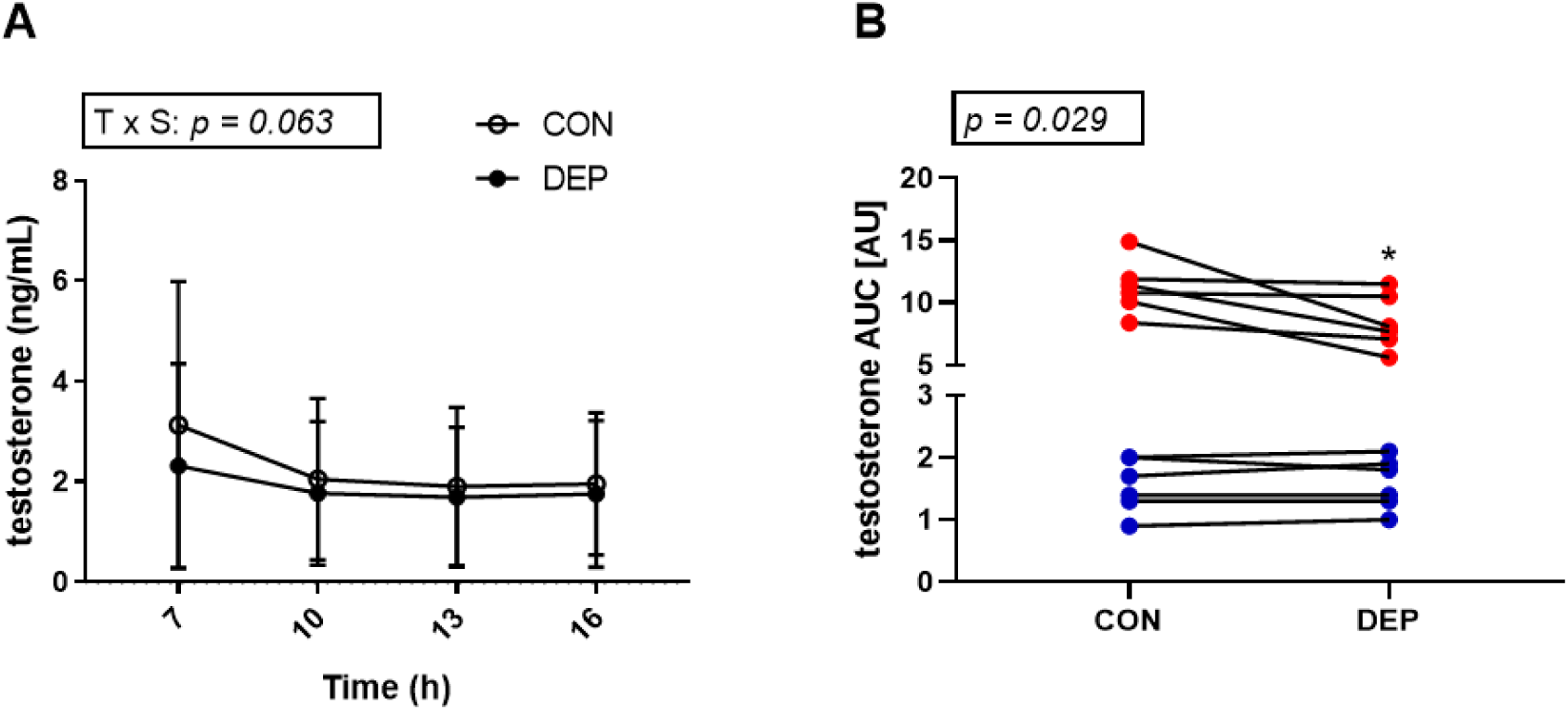
Plasma testosterone concentrations in control (CON) and sleep-deprived (DEP) conditions. Data are presented as mean ± SD (A). Area under the curve calculated for plasma cortisol concentrations. Red dots depict male subjects. Blue dots depict female subjects (B).

### Plasma cortisol levels

A significant interaction effect of sleep × time (*p=4.38E-5*) was observed for plasma cortisol levels. Consistent with the typical increase in cortisol observed during the later stages of sleep (47), plasma cortisol levels were significantly higher (*p=0.014*) in the CON condition than in the DEP condition at 0700 (wake time for the control condition). At 1000, plasma cortisol was similar in both sleep conditions (*p=0.940)*, but by 1600, cortisol was significantly higher in the DEP condition (*p=0.048)* (Figure 4A). Plasma cortisol area under the curve (1000-1600), was 21% higher during DEP than CON (*p=0.011)* (Figure 4B).

**Figure 4.**
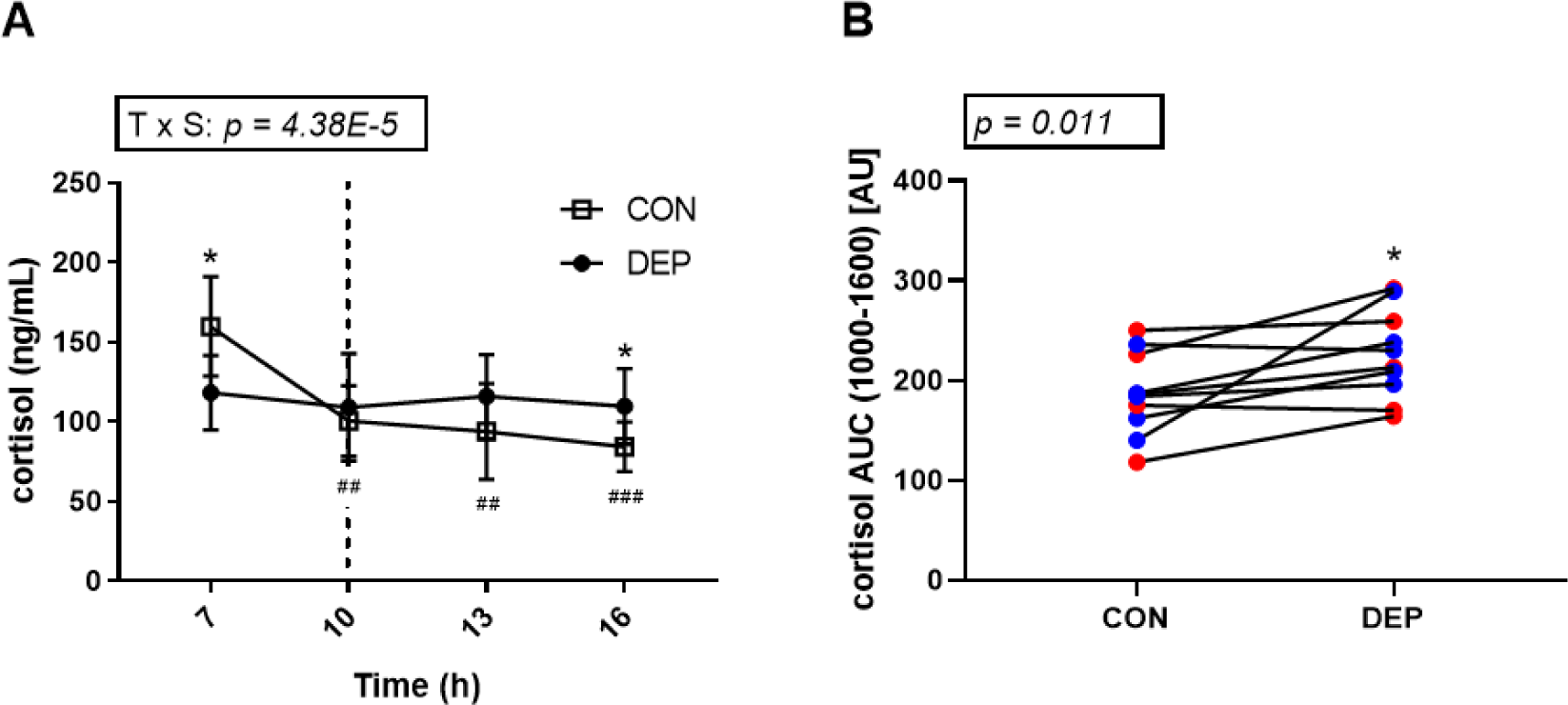
Plasma cortisol concentrations in control (CON) and sleep-deprived (DEP) conditions. ^*^; significantly different from the CON condition, *p<0.05*. ^##^; significantly different from the 700 timepoint in the CON group, *p<0.01.* ^###^; CON group was significantly different from the 700 timepoint in the CON group, *p<0.001.* Data are presented as mean ± SD (A). Area under the curve calculated for plasma cortisol concentrations from 1000 (dashed line, A) until the end of the protocol. ^*^; significantly different from the CON condition, *p<0.05*. Red dots depict male subjects. Blue dots depict female subjects (B).

### Insulin and IGF-1 levels

Plasma IGF-1 concentrations did not vary with time, sleep, or the combination of both (Figure 5A). Similarly, sleep deprivation did not influence the muscle expression levels of IGF1 mRNA isoforms IGF1-Ea and IGF1-Eb when measured in the post-prandial state (Figure 5B and 5C). Plasma insulin concentrations varied across the day, but there was no effect of sleep or the combination of sleep and time (Figure 5D).

**Figure 5.**
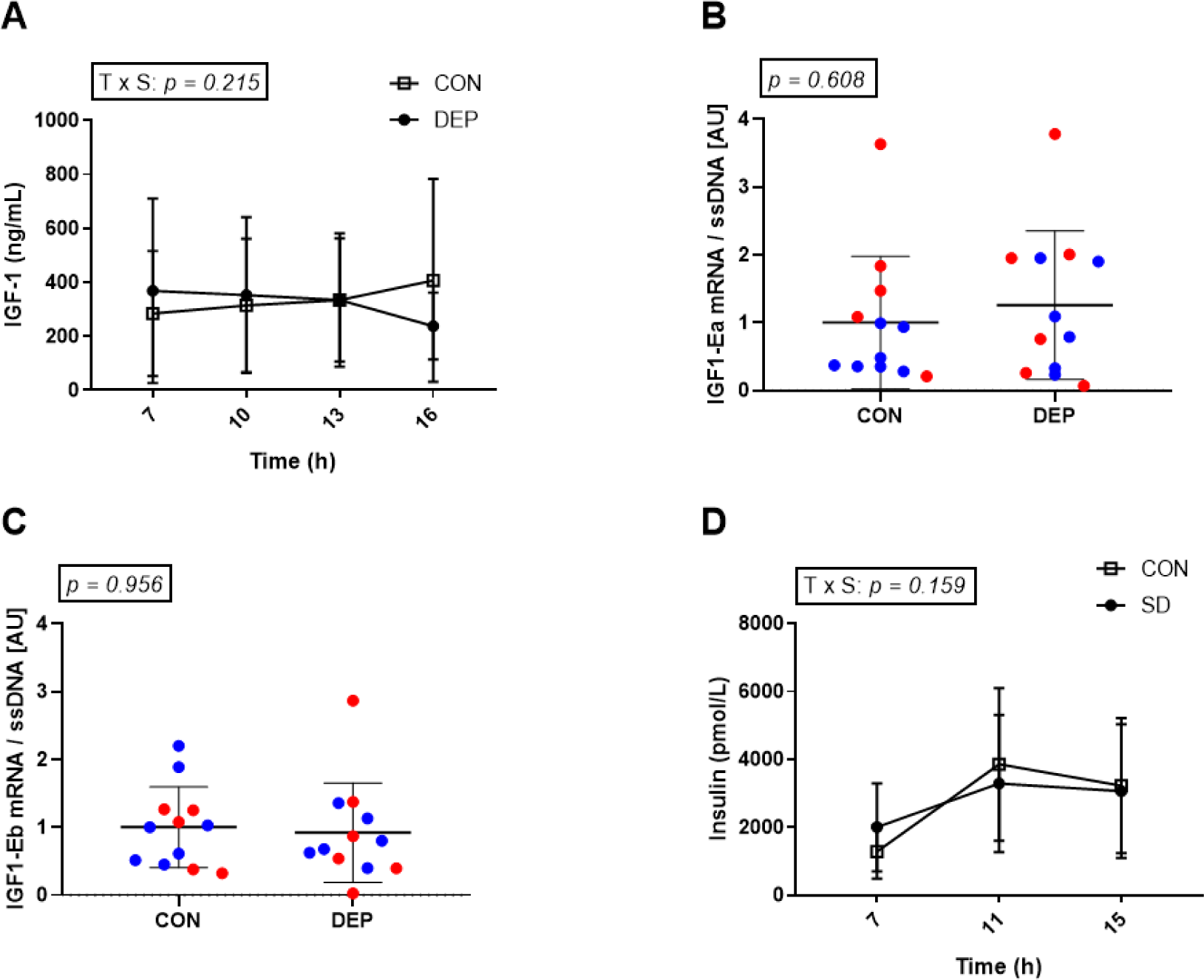
Plasma IGF-1 concentrations in control (CON) and sleep-deprived (DEP) conditions. (A). Muscle mRNA levels of the IGF-1 isoforms IGF1-Ea (B) and IGF1-Eb (C) in muscle biopsies collected at 1300. Red dots depict male subjects. Blue dots depict female subjects. Insulin concentrations in control (CON) and sleep-deprived (DEP) conditions (D). All data are presented as mean ± SD

### Gene expression

The muscle expression levels of core clock genes *ARNTL, CRY1* and *PER1* or muscle protein degradation markers *FBOX-32, MURF1, FOXO1* and *FOXO3* were assessed in muscle biopsies collected in the post-prandial state and did not change in response to sleep deprivation (Figure 6A–6G).

**Figure 6.**
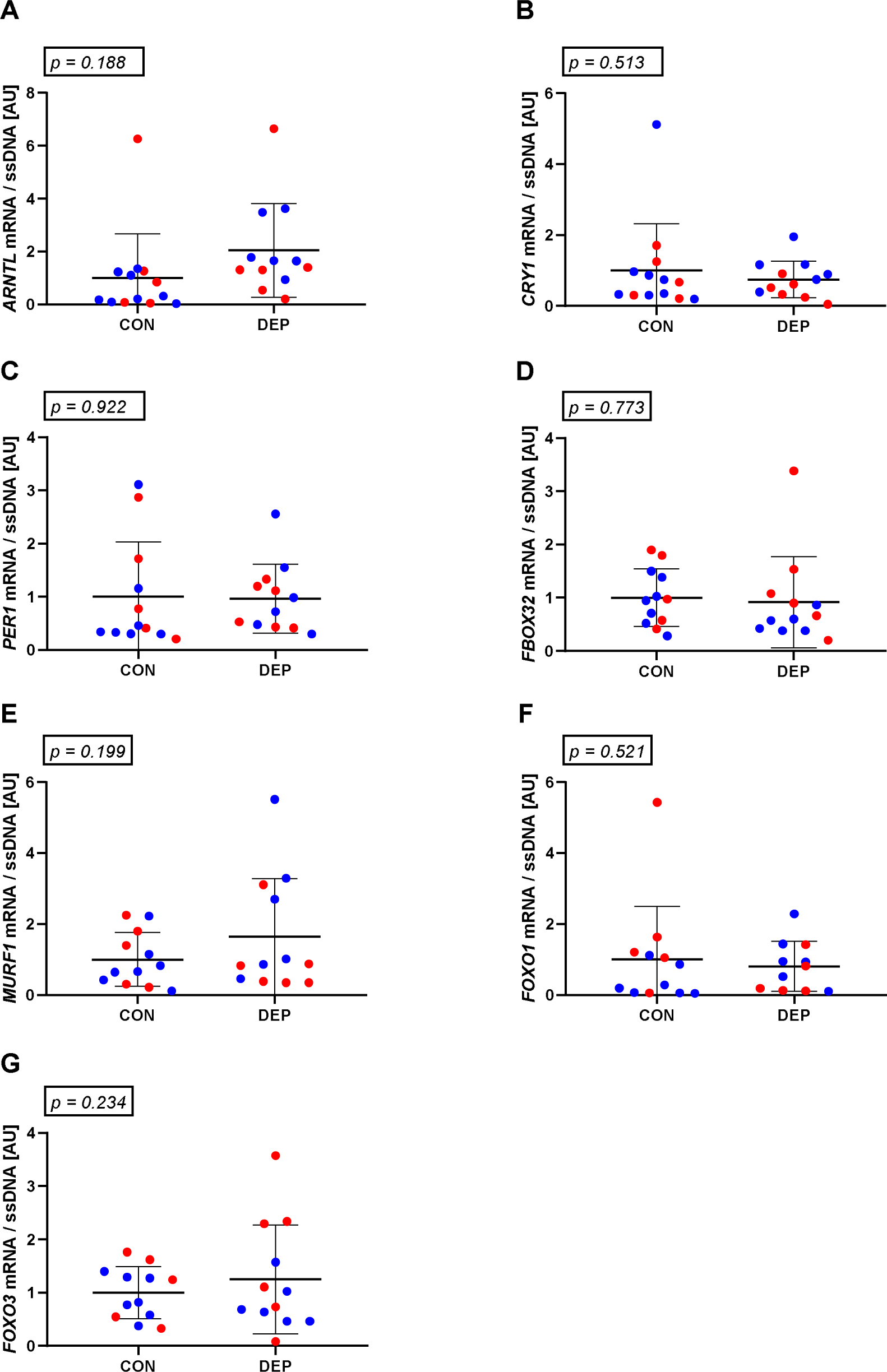
Muscle mRNA levels of *ARNTL* (A), *CRY1* (B), *PER1* (C), atrogin (*FBOX32*) (D), *MURF1* (E), *FOXO1* (F) and *FOXO3* (G) in muscle biopsies collected at 1300. Red dots depict male subjects. Blue dots depict female subjects. All data are presented as mean ± SD

## Discussion

Chronic sleep loss is a potent catabolic stressor (10, 42) that increases the risk of metabolic dysfunction (39) and is associated to a loss of muscle mass and function at the population level (31, 38). In this study, we have demonstrated that a single night of sleep deprivation is sufficient to induce anabolic resistance, reducing post-prandial skeletal muscle protein synthesis rates by 18%. This decrease was accompanied by an acute, pro-catabolic increase in plasma cortisol and a sex-specific reduction in plasma testosterone. Our study is the first to demonstrate that acute sleep deprivation blunts muscle protein synthesis, a key regulator of skeletal muscle turnover. It adds to early results reporting a reduction in muscle protein synthesis in chronic sleep restriction conditions (42) and provides insights into the mechanisms underlying the suppression of anabolism following an acute or chronic lack of sleep.

### Acute sleep deprivation decreases muscle protein synthesis

One night of sleep deprivation significantly reduced post-prandial skeletal muscle protein synthesis in a population of healthy young adults. In rodents, complete sleep deprivation is known to decrease muscle mass (13), muscle fibre cross-sectional area (15) and markers of the protein synthesis pathways. Only one study to date has investigated the effect of poor sleep on muscle protein synthesis in humans. Using a chronic sleep restriction model, Saner *et al.* recently reported that five consecutive nights of sleep restriction reduced muscle protein synthesis rates in healthy young males (42). Despite employing a different stable isotope (phenylalanine *versus* deuterated water), study design (acute cross-over design *versus* chronic parallel design) and population (males and females *versus* males only), our findings support those by Saner et al. Negative phenotypic outcomes associated with a period of chronic sleep deprivation likely reflect a metabolic shift towards catabolism due the accumulation of blunted anabolic responses to protein containing meals and physical activity. Our group has further discussed the results and implications of the Saner paper elsewhere (23).

A novel, exploratory outcome from our study highlights the potential for sex-specific responses to sleep deprivation. While the study was not powered to formally compare sexes, all of our male, but not female, participants experienced a numerical decrease in protein synthesis in the sleep-deprived *versus* control condition. Since we could not identify individual characteristics or behaviours consistent with the paradoxical increase in muscle protein synthesis observed in some of our female participants, our data may reflect a broader, sex-specific physiological response and warrants more focused attention.

To balance experimental rigor with subject comfort and improve the potential clinical translation of our data during both sleep trials, participants consumed a standardized meal prior to the first muscle biopsy. Dietary protein is a potent activator of muscle protein synthesis, potentially explaining the slightly higher protein synthesis rates we observed, compared to typical data obtained from fasted participants (27). By studying participants in the post-prandial state, we were able to conclude that acute sleep deprivation induces anabolic resistance, decreasing the capacity of muscle to respond to the typical anabolic stimulation triggered by dietary protein intake. These results have potential, far-reaching implications for the musculo-skeletal and metabolic health of populations including shiftworkers, new parents, students and older adults, who are at increased risk of acute and/or chronic sleep loss. Future research and clinical interventions prioritizing nutrition and/or protein-synthesis stimulating exercise (42) is warranted as these may represent practical and effective means of protecting muscle mass and function in sleep-deprived populations.

### Acute sleep deprivation promotes a less anabolic hormonal environemnt

Testosterone AUC was reduced by 22% following one night of sleep deprivation. Testosterone is the major androgenic hormone, but is also present in females, albeit in concentrations that are 10-fold lower than typical male levels (48). In our study, a sex-specific pattern was visible, where male, but not female, testosterone levels were attenuated by sleep deprivation. There is limited evidence describing how complete sleep deprivation alter testosterone daytime secretion patterns. In males, plasma testosterone fluctuates during the day, with concentrations increasing during sleep and gradually decreasing during waking periods (3, 45), with marginal circadian effects (3). A minimum of three hours of normal sleep, including paradoxical sleep opportunities (30), is required to increase testosterone. In an earlier study in healthy young men, one night of acute sleep deprivation did not alter 24 h testosterone AUC; however, a pattern similar to ours could be observed across the day (14). Collectively, these data first suggest that one night of sleep deprivation is sufficient to elicit a reduction in daytime testosterone concentrations. This effect appears particularly pronounced or inherent to males, where testosterone is a potent regulator of muscle protein synthesis both on the short (5 days) (43) and longer term (4 weeks) (46). However, acute exposure to testosterone is not sufficient to alter post-absorptive muscle protein synthesis or degradation rates over a 5-hour period (12). Whether transiently low testosterone levels can negatively impact muscle protein synthesis rates in the fasted and/or fed state is unknown and constitutes a challenge to validate experimentally. However, our results suggest that depressed testosterone secretion during the sleep deprivation period (30) is followed by another low testosterone secretion period during the daytime. Whether this phasic response contributes to anabolic resistance needs to be tested experimentally, but provides a possible mechanism for our observations.

### Acute sleep deprivation promotes a more catabolic hormonal environment but no difference in gene expression

Consistent with previous studies conducted in males, a cortisol response upon awakening was not observed following one night of acute sleep deprivation (47). This blunted cortisol response was accompanied by a chronically higher cortisol secretion across the day (10, 14). While our results are in line with these observations, the participants night-time calorie consumption needs to be acknowledged as a potential confounder. Previous studies have shown that over-night glucose infusion reduces cortisol levels (4). In our study, cortisol AUC was therefore calculated after the 1000 time point, after nutrient and energy intake was normalized. Further, post-hoc analyses revealed that, while the control group displayed a gradual, significant decrease in cortisol across the day, no differences were observed in the sleep deprived group at any time point, indicating a potential circadian misalignement. Cortisol has catabolic properties. In rats, corticosterone reduced muscle protein synthesis while increasing myofibrillar protein breakdown (21). In contrast, in humans, acute hypercortisolemia did not affect muscle fractional synthesis rates but blunted the net muscle protein balance (37), suggesting that cortisol preferentially increases muscle protein breakdown, rather than blunting muscle protein synthesis. Indeed, complete sleep deprivation can lead to a catabolic gene signature in human skeletal muscle (10), which might be reflective of an increase in muscle protein degradation. In our acute model, we however failed to observe any difference in the muscle expression levels of the proteolytic genes *FOXO1* and *FOXO3*, or in the expression levels of muscle specific atrogenes Atrogin-1 (*FBXO32*) and *MURF1*. This may be explained by the fact that our muscle biopsy was collected at a later time point (0730 *versus* 1300 in our study), but also by the post-prandial state of our participants at the time of sample collection. Indeed, consumption of a mixed meal attenuates ubiquitin-mediated proteolysis when compared to fasted (8). Whilst it should be kept in mind that acute studies essentialy report ‘snapshots of chronic processes, our choice to use a post prandial model has the advantage of providing a better reflection of time periods where the anabolic flux is greater. Supporting an effect of poor sleep on muscle protein degradation, some authors also recently suggested that poor sleep-induced hypercortisolemia might play a role in the development of sarcopenia (38), with potential sex-specific effects (6); however, further research is required to establish cause-and-effect relationships.

In contrast to others (9), we did not observe a decrease in the muscle expression levels of the core clock genes following a night of total sleep deprivation. Since the timing of muscle sample collection was different, it can be hypothesised that the muscle circadian rhythm might have been able to realign over this time period. It should also be acknowledged that night time calorie consumption constitutes a potential confounder as food intake can act as a “Zeitgeber” for peripheral tissues in mammals (11), including skeletal muscle. However, differences in core clock gene expression might not be reflective of a physiologically significant change. This warrants the comparison of more functional readouts, such as protein expression levels, which was not possible in this study due to tissue availability.

### Strenghts and Limitations

To improve compliance, comfort and retention, we were requested by our human ethics committee to allow participants to consume low-protein snacks (i.e. fruits and vegetable), and water *ad libitum* before normalizing calorie consumption at the 0700 time point, six hours before the first biopsy was collected. This strategy was effective in achieving similar plasma insulin levels across the two conditions at all time points. Using the home environment constituted a compromise as it avoids the need for habituation, which is a strength of this study, but rules out the ability to obtain overnight blood samples. Despite the presumed familiarity with their home sleep environment, our participants were mildly sleep-restricted in the week coming into both arms of the study, though total sleep times still fell within the stated sleep-wake time inclusion criterion. Sleep recorded on the pre-trial nights approximated their self-reported 7-h typical duration. Whether the mild-sleep restriction state had any impact on the results remains unclear. Future studies may consider the inclusion of laboratory-sleep trials and gold-standard polysomnographic sleep measures permitting analyses of sleep quality. While increasing participant burden, this would allow the characterisation of relationships between measures of sleep quality and physiological outcomes, including skeletal muscle protein synthesis and the hormonal environment.

Finally, in designing this clinical trial, we did not focus on sex-specific differences, nor did we power our study in order to detect such differences. Indeed, a wealth of literature indicate that muscle fractional synthesis rate, our primary outcome, is similar in males and females at rest and in response to anabolic stimulation (24, 49). However, the potential sex-based differences observed in this study prompt a dedicated investigation to better examine links between inadequate sleep and impaired skeletal muscle health in male and female cohorts.

## Acknowledgments

The authors wish to thank Dr Sarah Hall and Mr Teddy Ang for their excellent technical assistance.

## Author contributions

SL, GV and BA designed the study. SL, AM, OK, DC, SEA, AG collected the data. SL, EAL, OK, GV, SEA, DPJ processed and analysed the data. SL conducted statistical analyses and drafted the manuscript. SL and BA supervised the project. All authors commented on and edited the manuscript drafts.

## Additional Information

The authors have no conflict of interest to declare.

## References

1. Aisbett B, Condo D, Zacharewicz E, and Lamon S. The Impact of Shiftwork on Skeletal Muscle Health. Nutrients 9, 2017.

2. Ancoli-Israel S, Cole R, Alessi C, Chambers M, Moorcroft W, and Pollak CP. The role of actigraphy in the study of sleep and circadian rhythms. Sleep 26: 342–392, 2003.

3. Axelsson J, Ingre M, Akerstedt T, and Holmback U. Effects of acutely displaced sleep on testosterone. J Clin Endocrinol Metab 90: 4530–4535, 2005.

4. Benedict C, Kern W, Schmid SM, Schultes B, Born J, and Hallschmid M. Early morning rise in hypothalamic-pituitary-adrenal activity: a role for maintaining the brain's energy balance. Psychoneuroendocrinology 34: 455–462, 2009.

5. Bescos R, Boden MJ, Jackson ML, Trewin AJ, Marin EC, Levinger I, Garnham A, Hiam DS, Falcao-Tebas F, Conte F, Owens JA, Kennaway DJ, and McConell GK. Four days of simulated shift work reduces insulin sensitivity in humans. Acta physiologica (Oxford, England) 223: e13039, 2018.

6. Buchmann N, Spira D, Norman K, Demuth I, Eckardt R, and Steinhagen-Thiessen E. Sleep, Muscle Mass and Muscle Function in Older People. Dtsch Arztebl Int 113: 253–260, 2016.

7. Buxton OM, Pavlova M, Reid EW, Wang W, Simonson DC, and Adler GK. Sleep restriction for 1 week reduces insulin sensitivity in healthy men. Diabetes 59: 2126–2133, 2010.

8. Carbone JW, Margolis LM, McClung JP, Cao JJ, Murphy NE, Sauter ER, Combs GF, Jr., Young AJ, and Pasiakos SM. Effects of energy deficit, dietary protein, and feeding on intracellular regulators of skeletal muscle proteolysis. Faseb j 27: 5104–5111, 2013.

9. Cedernaes J, Osler ME, Voisin S, Broman JE, Vogel H, Dickson SL, Zierath JR, Schioth HB, and Benedict C. Acute Sleep Loss Induces Tissue-Specific Epigenetic and Transcriptional Alterations to Circadian Clock Genes in Men. J Clin Endocrinol Metab 100: E1255–1261, 2015.

10. Cedernaes J, Schonke M, Westholm JO, Mi J, Chibalin A, Voisin S, Osler M, Vogel H, Hornaeus K, Dickson SL, Lind SB, Bergquist J, Schioth HB, Zierath JR, and Benedict C. Acute sleep loss results in tissue-specific alterations in genome-wide DNA methylation state and metabolic fuel utilization in humans. Science advances 4: eaar8590, 2018.

11. Challet E. The circadian regulation of food intake. Nat Rev Endocrinol 15: 393–405, 2019.

12. Church DD, Pasiakos SM, Wolfe RR, and Ferrando AA. Acute testosterone administration does not affect muscle anabolism. Nutrition & metabolism 16: 56, 2019.

13. Dattilo M, Antunes HK, Medeiros A, Monico-Neto M, Souza Hde S, Lee KS, Tufik S, and de Mello MT. Paradoxical sleep deprivation induces muscle atrophy. Muscle Nerve 45: 431–433, 2012.

14. Dattilo M, Antunes HKM, Galbes NMN, Monico-Neto M, H Dess, Dos Santos Quaresma MVL, Lee KS, Ugrinowitsch C, Tufik S, and MT DEM. Effects of Sleep Deprivation on Acute Skeletal Muscle Recovery after Exercise. Med Sci Sports Exerc 52: 507–514, 2020.

15. de Sa Souza H, Antunes HK, Dattilo M, Lee KS, Monico-Neto M, de Campos Giampa SQ, Phillips SM, Tufik S, and de Mello MT. Leucine supplementation is anti-atrophic during paradoxical sleep deprivation in rats. Amino Acids 48: 949–957, 2016.

16. English KL, Mettler JA, Ellison JB, Mamerow MM, Arentson-Lantz E, Pattarini JM, Ploutz-Snyder R, Sheffield-Moore M, and Paddon-Jones D. Leucine partially protects muscle mass and function during bed rest in middle-aged adults. Am J Clin Nutr 103: 465–473, 2016.

17. Glynn EL, Fry CS, Timmerman KL, Drummond MJ, Volpi E, and Rasmussen BB. Addition of carbohydrate or alanine to an essential amino acid mixture does not enhance human skeletal muscle protein anabolism. The Journal of nutrition 143: 307–314, 2013.

18. Hamer HM, Wall BT, Kiskini A, de Lange A, Groen BB, Bakker JA, Gijsen AP, Verdijk LB, and van Loon LJ. Carbohydrate co-ingestion with protein does not further augment post-prandial muscle protein accretion in older men. Nutrition & metabolism 10: 15, 2013.

19. Harfmann BD, Schroder EA, and Esser KA. Circadian rhythms, the molecular clock, and skeletal muscle. Journal of biological rhythms 30: 84–94, 2015.

20. Horne JA and Ostberg O. A self-assessment questionnaire to determine morningness-eveningness in human circadian rhythms. Int J Chronobiol 4: 97–110, 1976.

21. Kayali AG, Young VR, and Goodman MN. Sensitivity of myofibrillar proteins to glucocorticoid-induced muscle proteolysis. Am J Physiol 252: E621–626, 1987.

22. Kecklund G and Axelsson J. Health consequences of shift work and insufficient sleep. BMJ 355: i5210, 2016.

23. Knowles OE. No time to sleep on it - start exercising! J Physiol 598: 2059–2060, 2020.

24. Knowles OE, Aisbett B, Main LC, Drinkwater EJ, Orellana L, and Lamon S. Resistance Training and Skeletal Muscle Protein Metabolism in Eumenorrheic Females: Implications for Researchers and Practitioners. Sports Med, 2019.

25. Koopman R, Beelen M, Stellingwerff T, Pennings B, Saris WH, Kies AK, Kuipers H, and van Loon LJ. Coingestion of carbohydrate with protein does not further augment postexercise muscle protein synthesis. American journal of physiology Endocrinology and metabolism 293: E833–842, 2007.

26. Kowall B, Lehnich AT, Strucksberg KH, Fuhrer D, Erbel R, Jankovic N, Moebus S, Jockel KH, and Stang A. Associations among sleep disturbances, nocturnal sleep duration, daytime napping, and incident prediabetes and type 2 diabetes: the Heinz Nixdorf Recall Study. Sleep Med 21: 35–41, 2016.

27. Kumar V, Selby A, Rankin D, Patel R, Atherton P, Hildebrandt W, Williams J, Smith K, Seynnes O, Hiscock N, and Rennie MJ. Age-related differences in the dose-response relationship of muscle protein synthesis to resistance exercise in young and old men. J Physiol 587: 211–217, 2009.

28. Lamon S, Zacharewicz E, Arentson-Lantz E, Gatta PA, Ghobrial L, Gerlinger-Romero F, Garnham A, Paddon-Jones D, and Russell AP. Erythropoietin Does Not Enhance Skeletal Muscle Protein Synthesis Following Exercise in Young and Older Adults. Front Physiol 7: 292, 2016.

29. Leproult R and Van Cauter E. Effect of 1 week of sleep restriction on testosterone levels in young healthy men. JAMA 305: 2173–2174, 2011.

30. Luboshitzky R, Zabari Z, Shen-Orr Z, Herer P, and Lavie P. Disruption of the nocturnal testosterone rhythm by sleep fragmentation in normal men. J Clin Endocrinol Metab 86: 1134–1139, 2001.

31. Lucassen EA, de Mutsert R, le Cessie S, Appelman-Dijkstra NM, Rosendaal FR, van Heemst D, den Heijer M, Biermasz NR, and group NEOs. Poor sleep quality and later sleep timing are risk factors for osteopenia and sarcopenia in middle-aged men and women: The NEO study. PLoS One 12: e0176685, 2017.

32. Lundby C, Nordsborg N, Kusuhara K, Kristensen KM, Neufer PD, and Pilegaard H. Gene expression in human skeletal muscle: alternative normalization method and effect of repeated biopsies. Eur J Appl Physiol 95: 351–360, 2005.

33. Mamerow MM, Mettler JA, English KL, Casperson SL, Arentson-Lantz E, Sheffield-Moore M, Layman DK, and Paddon-Jones D. Dietary protein distribution positively influences 24-h muscle protein synthesis in healthy adults. The Journal of nutrition 144: 876–880, 2014.

34. Miller BF, Hansen M, Olesen JL, Flyvbjerg A, Schwarz P, Babraj JA, Smith K, Rennie MJ, and Kjaer M. No effect of menstrual cycle on myofibrillar and connective tissue protein synthesis in contracting skeletal muscle. American journal of physiology Endocrinology and metabolism 290: E163–e168, 2006.

35. Monico-Neto M, Antunes HK, Dattilo M, Medeiros A, Souza HS, Lee KS, de Melo CM, Tufik S, and de Mello MT. Resistance exercise: a non-pharmacological strategy to minimize or reverse sleep deprivation-induced muscle atrophy. Medical hypotheses 80: 701–705, 2013.

36. Paddon-Jones D, Sheffield-Moore M, Cree MG, Hewlings SJ, Aarsland A, Wolfe RR, and Ferrando AA. Atrophy and impaired muscle protein synthesis during prolonged inactivity and stress. J Clin Endocrinol Metab 91: 4836–4841, 2006.

37. Paddon-Jones D, Sheffield-Moore M, Creson DL, Sanford AP, Wolf SE, Wolfe RR, and Ferrando AA. Hypercortisolemia alters muscle protein anabolism following ingestion of essential amino acids. American journal of physiology Endocrinology and metabolism 284: E946–953, 2003.

38. Piovezan RD, Abucham J, Dos Santos RV, Mello MT, Tufik S, and Poyares D. The impact of sleep on age-related sarcopenia: Possible connections and clinical implications. Ageing Res Rev 23: 210–220, 2015.

39. Reutrakul S and Van Cauter E. Sleep influences on obesity, insulin resistance, and risk of type 2 diabetes. Metabolism 84: 56–66, 2018.

40. Reynolds AC, Dorrian J, Liu PY, Van Dongen HP, Wittert GA, Harmer LJ, and Banks S. Impact of five nights of sleep restriction on glucose metabolism, leptin and testosterone in young adult men. PLoS One 7: e41218, 2012.

41. Russell AP. Molecular regulation of skeletal muscle mass. Clinical and experimental pharmacology & physiology 37: 378–384, 2010.

42. Saner NJ, Lee MJ, Pitchford NW, Kuang J, Roach GD, Garnham A, Stokes T, Phillips SM, Bishop DJ, and Bartlett JD. The effect of sleep restriction, with or without high-intensity interval exercise, on myofibrillar protein synthesis in healthy young men. J Physiol, 2020.

43. Sheffield-Moore M, Urban RJ, Wolf SE, Jiang J, Catlin DH, Herndon DN, Wolfe RR, and Ferrando AA. Short-term oxandrolone administration stimulates net muscle protein synthesis in young men. J Clin Endocrinol Metab 84: 2705–2711, 1999.

44. Staples AW, Burd NA, West DW, Currie KD, Atherton PJ, Moore DR, Rennie MJ, Macdonald MJ, Baker SK, and Phillips SM. Carbohydrate does not augment exercise-induced protein accretion versus protein alone. Med Sci Sports Exerc 43: 1154–1161, 2011.

45. Touitou Y, Motohashi Y, Reinberg A, Touitou C, Bourdeleau P, Bogdan A, and Auzeby A. Effect of shift work on the night-time secretory patterns of melatonin, prolactin, cortisol and testosterone. Eur J Appl Physiol Occup Physiol 60: 288–292, 1990.

46. Urban RJ, Bodenburg YH, Gilkison C, Foxworth J, Coggan AR, Wolfe RR, and Ferrando A. Testosterone administration to elderly men increases skeletal muscle strength and protein synthesis. Am J Physiol 269: E820–826, 1995.

47. Vargas I and Lopez-Duran N. The cortisol awakening response after sleep deprivation: Is the cortisol awakening response a "response" to awakening or a circadian process? Journal of health psychology: 1359105317738323, 2017.

48. Vingren JL, Kraemer WJ, Ratamess NA, Anderson JM, Volek JS, and Maresh CM. Testosterone Physiology in Resistance Exercise and Training. Sports Medicine 40: 1037–1053, 2010.

49. West DW, Burd NA, Churchward-Venne TA, Camera DM, Mitchell CJ, Baker SK, Hawley JA, Coffey VG, and Phillips SM. Sex-based comparisons of myofibrillar protein synthesis after resistance exercise in the fed state. J Appl Physiol (1985) 112: 1805–1813, 2012.

50. Wu Y, Zhai L, and Zhang D. Sleep duration and obesity among adults: a meta-analysis of prospective studies. Sleep Med 15: 1456–1462, 2014.

51. Zhao W, Pan J, Wang X, Wu Y, Bauman WA, and Cardozo CP. Expression of the muscle atrophy factor muscle atrophy F-box is suppressed by testosterone. Endocrinology 149: 5449–5460, 2008.

